# Characterization of coral-associated microbial aggregates (CAMAs) within tissues of the coral *Acropora hyacinthus*

**DOI:** 10.1101/576488

**Authors:** Naohisa Wada, Mizuki Ishimochi, Taeko Matsui, F. Joseph Pollock, Sen-Lin Tang, Tracy D. Ainsworth, Bette L. Willis, Nobuhiro Mano, David G. Bourne

## Abstract

Bacterial diversity associated with corals has been studied extensively, however, localization of bacterial associations within the holobiont is still poorly resolved. Here we provide novel insight into the localization of coral-associated microbial aggregates (CAMAs) within tissues of the coral *Acropora hyacinthus*. In total, 318 and 308 CAMAs were characterized via histological and fluorescent *in situ* hybridization (FISH) approaches respectively, and shown to be distributed extensively throughout coral tissues collected from five sites in Japan and Australia. The densities of CAMAs within the tissues were negatively correlated with the distance from the coastline (i.e. lowest densities at offshore sites). CAMAs were randomly distributed across the six coral tissue regions investigated. Within each CAMA, bacterial cells had similar morphological characteristics, but bacterial morphologies varied among CAMAs, with at least five distinct types identified. Identifying the location of microorganisms associated with the coral host is a prerequisite for understanding their contributions to fitness. Localization of tissue-specific communities housed within CAMAs is particularly important, as these communities are potentially important contributors to vital metabolic functions of the holobiont.

## Introduction

Scleractinian corals associate with broad consortium of microorganisms, including endosymbiont dinoflagellates (Symbiodiniaceae), protozoa, fungi, bacteria, archaea and viruses, which collectively are termed the coral holobiont^1–3^. The importance of symbiotic dinoflagellates in provisioning the coral host with essential nutrients through translocated photosynthates has been established^e.g^.^4,5^, however the roles of other microorganisms within the holobiont are less well understood (reviewed in^3^). Some of the functions attributed to coral-associated microbiota include supply of essential nutrients and vitamins through processes such as nitrogen fixation^6–9^ and metabolizing dimethylsulfoniopropionate (DMSP) to produce biologically important byproducts like dimethylsulfide^10^. The coral microbiota is also likely important for directly facilitating disease resistance through production of antimicrobials^11,12^ or indirectly by preventing colonization of opportunistic or pathogenic organisms^13,14^.

Corals are considered simple metazoans, but despite their basal phylogenetic position, they nevertheless form complex three-dimensional structures. Anatomically, the coral consists of (1) a surface mucus layer, (2) polyps consisting of feeding tentacles, actinopharynx, mesenteries (including their filaments), and walls (surface and basal), (3) a gastrovascular system that includes the gastrovascular cavity (formerly coelenteron) and connecting canals and (4) an external calcium carbonate skeleton^15.72^. The coral tissue layers are composed of epithelia, calicodermis (formerly calicoblastic epithelium) and the gastrodermis, which also contains the Symbiodinaceae. Within all these microhabitat niches, bacteria can reside as either transient communities or established symbionts with functional roles that may be positive, neutral or negative to the coral holobiont^16^. A multitude of studies have reported on the diversity of the coral-associated microbial communities, in some cases finding conserved microbial communities associated with some coral species, and in others finding shifting microbial profiles that reflect varying geographic, temporal or health status patterns^17–20^. To understand the significance of coral-microbial associations, care must be exercised so that diversity patterns reflect the specific ecological niche that these communities inhabit, such as the surface mucus layer, tissue layers, and/or the skeleton^21–28^. Defining the locations of specific microorganisms is essential for elucidating the importance of their role within the holobiont. For example, mucus bacteria are more likely to have a loose association with the coral host, being sloughed off as the mucus is exuded from the corals^29^. Conversely tissue-associated microorganisms are potentially more integrated in shared metabolic pathways and may reside in specific associations with host coral as a consequence of potential host selection^6,20,30^.

To date, few studies have precisely localized bacterial communities within coral tissues. Studies that have focused on localization often find that bacterial communities within coral cell layers (i.e., epidermal and gastrodermal epithelia) form aggregations termed coral-associated microbial aggregates (CAMAs)^31–34^. CAMAs were first reported as potential pathogens when observed within healthy tissues of Caribbean corals displaying signs of white-band disease^35,36^. Further studies subsequently reported that CAMAs are widespread in tissues of healthy corals sampled from geographically dispersed areas^32,36,37^. To date, CAMAs have been reported from 5 species of corals in the Caribbean^35,36^ and 24 species from the Indo-Pacific^32^, although their frequency varies among coral genera, with the genera *Acropora, Porites*, and *Pocillopora* most commonly hosting bacterial aggregates^32^. Identification of the microorganisms that constitute these CAMAs has been poorly resolved. Neave et al. (2016) visualized aggregates (i.e., “cyst-like aggregations”) of *Endozoicomonas* within tissues of *Stylophora pistillata* at the interface of the epidermis and gastrodermis and confirmed their distribution in samples taken from widely-separated biogeographic regions^38,39^. However, the fine-scale spatial distributions of microorganisms and potential microhabitat-associated structure of the microbiome still have not been clarified. To facilitate a more comprehensive understanding of the roles bacterial communities may play in the coral holobiont, an improved understanding of the localization of these microbial communities is essential, particularly tissue-associated bacterial communities housed within structures termed coral-associated microbial aggregates (CAMAs). Here, we visualize the localization, distribution and morphology of CAMAs associated with the coral *Acropora hyacinthus* sampled from Sesoko Island, Okinawa, Japan and sites in the northern and central regions of the Great Barrier Reef (GBR) Australia, located along an inshore to offshore gradient.

## Results and Discussion

### Comparison of CAMA prevalence among geographic locations

At the time of field collection, all 48 colonies of *A. hyacinthus* sampled (one fragment from each colony), from the 5 geographic locations (see **Fig. 1**), appeared visually healthy. This was confirmed by subsequent histological analyses, which found that all tissues displayed normal cell morphology, including no signs of fragmentation, wound repair or necrosis (as per criteria in^40^). In total, 318 CAMAs were characterized via histology (**Fig. 2a**) with CAMAs detected in fragments from 27 of the 48 sampled colonies (~56%). The vast majority stained basophilic (95.9%), compared to only 4.1% staining eosinophilic with the hematoxylin and eosin staining procedure. At the Sesoko Island site, tissues derived from the fragments of all 10 sampled colonies contained CAMAs (**Fig. 2c**), whereas at the other four sites, CAMAs were observed only in tissues of some of the coral fragments. For example, CAMAs were clearly visible in 80% of Inner Shelf samples, 30% of Lizard Island, 25% of Outer Shelf, and 40% of Orpheus Island samples (n=10 samples at all sites, except at the Outer Shelf site where n=8 samples). In general, the prevalence of CAMAs was significantly higher at the Sesoko Island site than at the Lizard Island, Outer Shelf, and Orpheus Island sites (Fisher’s exact test: *p* = 0.00099, **Fig. 2c**, and see more detail of the pairwise test in **Suppl. table S1**).

**Fig. 1.**
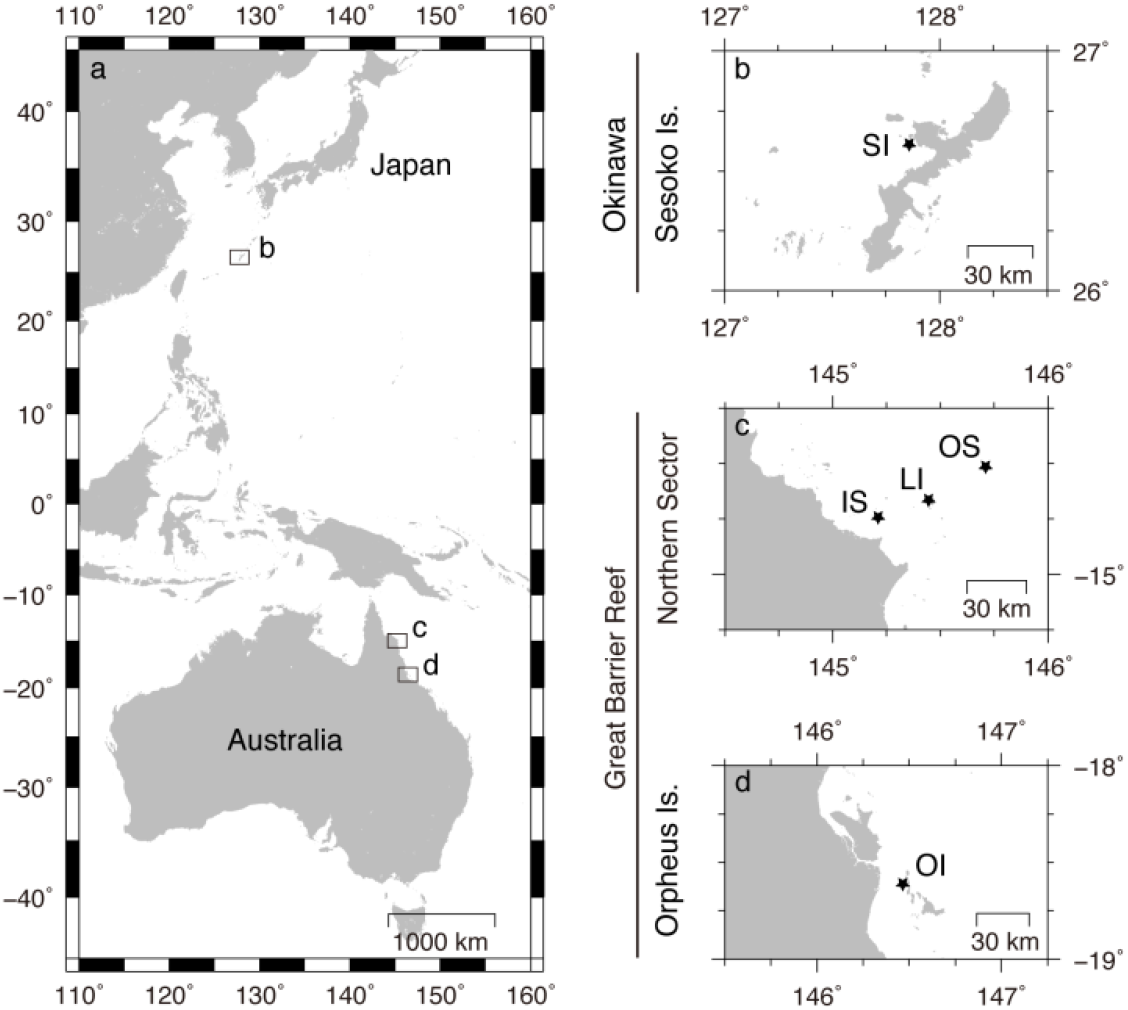
Map showing the locations of five study sites in two countries: **(a, b)** Sesoko Island (SI) in Okinawa, Japan; **(a, c)** Inner Shelf (IS), Lizard Island (LI) and Outer Shelf (OS) sites in the Northern Great Barrier Reef; and **(a, d)** Orpheus Island (OI) in the central Great Barrier Reef, Australia.

**Fig. 2.**
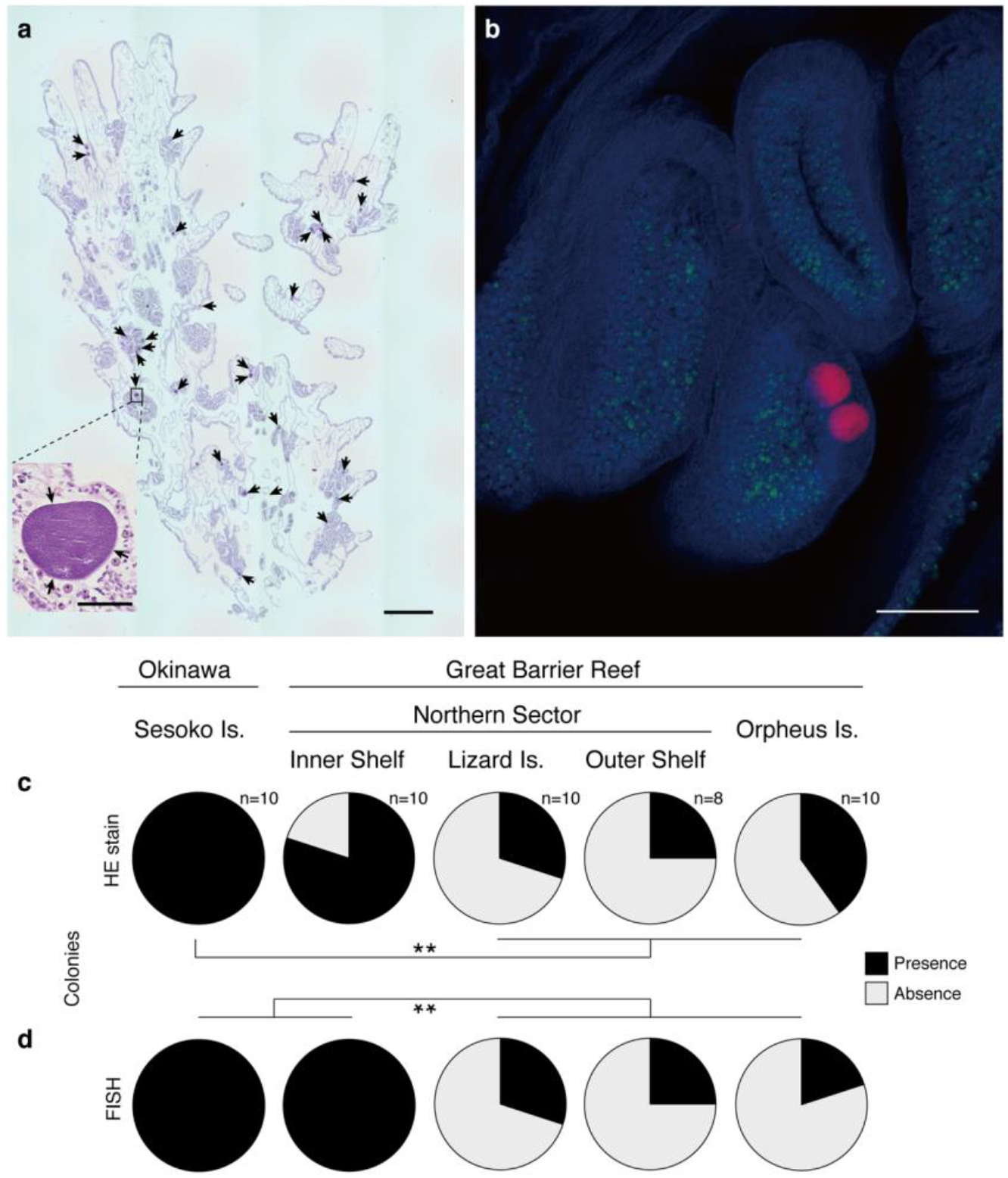
Appearance, occurrence of CAMAs in the coral *Acropora hyacinthus*. **(a)** Numerous CAMAs (indicated by arrows) are visible in a histological section stained by hematoxylin and eosin of a colony from Sesoko, Japan. In the panel, small photo shows close-up of a CAMA located in a mesentery of a polyp. **(b)** Right panel shows two of CAMAs (red) distributed with Symbiodiniaceae (green) in the tentacle (coral tissue: blue) of a sample from the Inner Shelf, GBR using FISH. **(c, d)** Pie diagrams showing the proportion of colonies sampled that contained CAMAs at five sites in Japan and Australia based on the detections of **(c)** HE stain and **(d)** FISH. **(c)** A significantly higher proportion of Sesoko Is. samples contained CAMAs (including basophilic and eosinophilic) than samples collected from other sites (Lizard Is., Outer Shelf northern GBR site, and Orpheus Is.). **(d)** In the FISH experiment, both proportions of CAMA in samples from Sesoko Is. and Inner Shelf were also significantly higher than in samples from other three sites (** *p*<0001; Fisher’s exact test followed by a Benjamini-Hochberg false discovery rate correction). Scale bars indicate 600 μm **(a)**, 50 μm (**small panel in a**) and 100 μm **(b)**.

A total of 308 CAMAs were also identified by FISH in the serial-sectioned tissues derived from the sampled colonies (again observed in 27 fragments from 48 sampled colonies) (**Fig. 2b**). CAMAs were observed from all samples derived from Sesoko Island and the Inner Shelf (**Fig. 2d**). In contrast CAMAs were observed in 30% of Lizard Island, 25% of Outer Shelf, and 20% of Orpheus Island samples. The prevalence of CAMAs detected by FISH was significantly higher at both Sesoko Island and Inner Shelf than at the other sites (Lizard Island, Outer Shelf, and Orpheus Island sites; Fisher’s exact test: *p* = 0.0000019, **Fig. 2d** and see more detail of the pairwise test in **Suppl. table S1**).

The density of basophilic (i.e. stained with hematoxylin) CAMAs in tissues was highest in the Sesoko Island samples (n=10) with 20.13 ± 17.1 per cm^2^ (average and S.E.), compared to 6.78 ± 9.3 per cm^2^ for the Inner Shelf (n=8) samples, 0.92 ± 0.1 per cm^2^ at Lizard Island (n=3) and 3.55 ± 3.3 per cm^2^ in Orpheus Island (n=3) samples (see more detail in **Suppl. table S2**). Basophilic CAMAs were not detected in samples of the Outer Shelf corals. Eosinophilic (i.e., stained by eosin) CAMAs were only detected in coral fragments derived from three sites, with 2.32, 5.39 ± 6.7, and 1.28 ± 2.8 per cm^2^ in Inner Shelf (n=1), Outer Shelf (n=3) and Orpheus Island (n=2) samples, respectively (see more detail in **Suppl. table S2**). The densities of CAMAs detected by FISH were reflective of the histological patterns with 18.72 ± 12.7 per cm^2^ in Sesoko Island (n=10), 7.90 ± 8.2 per cm^2^ in Inner Shelf (n=10), 1.26 ± 0.6 per cm^2^ in Lizard Island (n=3), 0.90 ± 0.5 per cm^2^ in Outer Shelf (n=2), and 4.48 ± 4.2 per cm^2^ in Orpheus Island (n=2) samples (see more detail in **Suppl. table S2**). In one sample from Sesoko Island, 48.9 basophilic CAMAs were detected per cm^2^ of tissue (41.1 detected by FISH in the serial section), the greatest density of CAMAs observed across all 48 samples investigated. The maximum number observed in a sample from the Outer Shelf region was 10.10 eosinophilic CAMAs per cm^2^.

The patterns of CAMAs prevalence derived from histological and FISH analysis confirm that CAMAs commonly occur in healthy tissues of the coral *Acropora hyacinthus* collected from sites in Japan and Australia separated by more than 40 degrees of latitude. Although CAMAs were not detected in all tissue samples collected, this may reflect limitations in the area of coral tissue that can be surveyed via histological approaches. The presence of CAMAs in a histological section will depend on the tissue sectioned (i.e., the location of the fragment on the colony and on the section from that fragment), and on the scale and orientation of the section. Indeed, our results showed variation of both presence and density of CAMAs, even from samples derived from the same site (see more detail in **Suppl. table. S2**). Therefore, although not all coral tissue samples (and therefore not all colonies) were found to host CAMAs, we cannot exclude the possibility that other tissue areas of the same colony had CAMAs present. Other studies have also reported that CAMAs are common in tissues of many coral species in the Caribbean^35,36^. Indo-Pacific and Red Sea^32,33,37,38^. In particular, CAMAs were common in species of *Acropora, Porites*, and *Pocillopora*, although often their presence was patchy within a population sample^32^.

Interestingly the distance from the coast negatively correlated with the proportion and density of CAMAs within coral tissues across the sampling sites (**Table 1**). A predictive model was constructed which yielded an ordinary negative binomial regression which was found preferable to a zero-inflated model (Vuong test; Voung z-statistic= 22.57413, *p*-value < 0.001 in the basophilic CAMAs, and Voung z-statistic= 3.120e-6, *p*<0.001 in CAMAs detected by FISH). Therefore an increased distance from the coastline influenced negatively the proportion and density of both basophilic CAMAs and CAMAs detected by FISH (**Fig. 3a, c**). The density of eosinophilic CAMAs did not follow the negative binominal regression model (**Table 1, Fig. 3b**). Although our study provides only a snapshot of five sites sampled at one time point, it suggests that inshore reef environments may promote the development of CAMAs within coral tissues. The inshore site at Sesoko Island has high nutrient influxes, especially phosphates^41,42^. Similarly, nearshore GBR sites are influenced by influxes of dissolved nutrients from terrestrial runoff, with higher concentrations typically found at inshore compared with offshore sites^43^. Temperature fluctuations may also influence CAMA abundance, although differences in seasonal temperature fluctuations are minimal across the northern GBR sites^44^. The coral microbiome community has been shown to shift in response to environmental stressors^29,45–49^, thus water quality parameters influencing microbiological composition and function^50^ could stimulate CAMA abundance. Further studies, particularly of potential links between nutrient levels and CAMA development, are needed to understand what might drive the increased prevalence and density of CAMAs in coral tissues at inshore sites, and to determine if hosting more CAMAs is beneficial to corals or is an indicator of negative impacts on the coral holobiont.

**Table 1.**
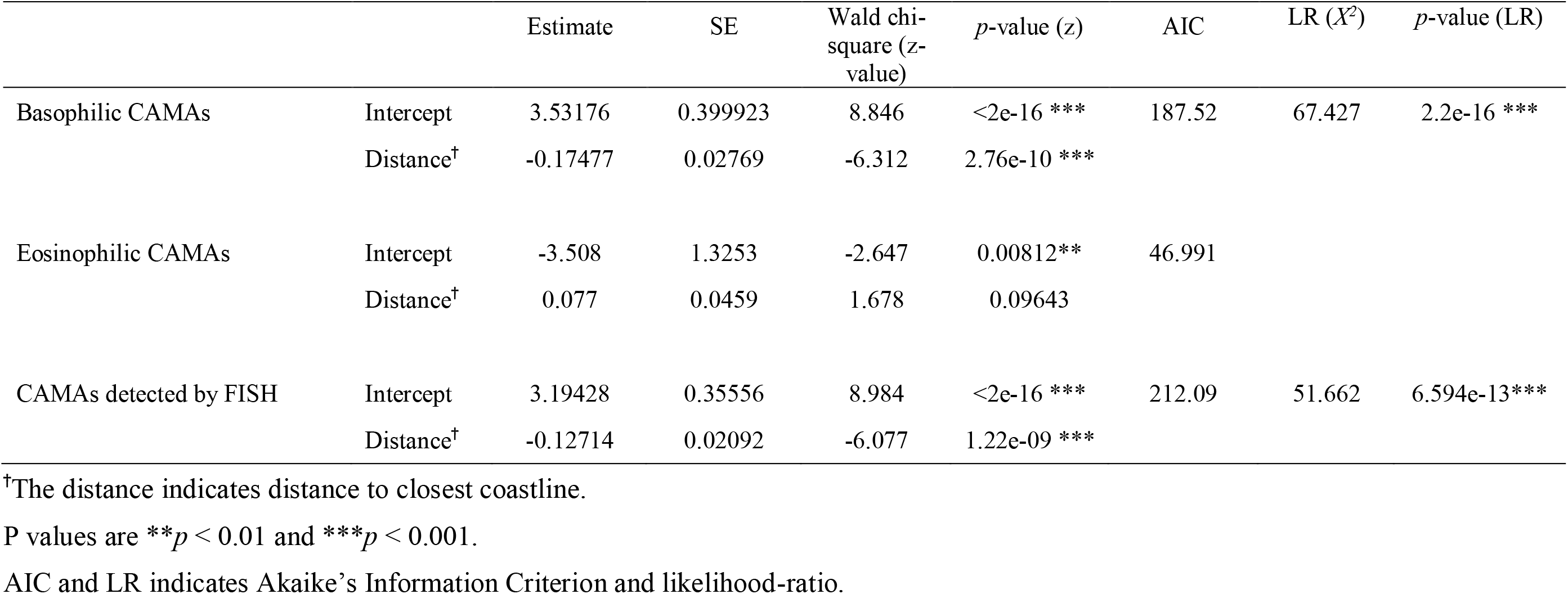
Results from the generalized linear regression models with negative binominal distribution testing the effect of the distance from closest coastline on the densities of CAMAs

| | | Estimate | SE | Wald chi-square (z-value) | *p*-value (z) | AIC | LR ($X^2$) | *p*-value (LR) |
| --- | --- | --- | --- | --- | --- | --- | --- | --- |
| Basophilic CAMAs | Intercept | 3.53176 | 0.399923 | 8.846 | <2e-16 *** | 187.52 | 67.427 | 2.2e-16 *** |
|  | Distance<sup>†</sup> | -0.17477 | 0.02769 | -6.312 | 2.76e-10 *** |  |  |  |
| Eosinophilic CAMAs | Intercept | -3.508 | 1.3253 | -2.647 | 0.00812** | 46.991 |  |  |
|  | Distance<sup>†</sup> | 0.077 | 0.0459 | 1.678 | 0.09643 |  |  |  |
| CAMAs detected by FISH | Intercept | 3.19428 | 0.35556 | 8.984 | <2e-16 *** | 212.09 | 51.662 | 6.594e-13*** |
|  | Distance<sup>†</sup> | -0.12714 | 0.02092 | -6.077 | 1.22e-09 *** |  |  |  |

<sup>†</sup>The distance indicates distance to closest coastline.
P values are \*\* $p < 0.01$ and \*\*\* $p < 0.001$ .
AIC and LR indicates Akaike's Information Criterion and likelihood-ratio.

**Fig. 3.**
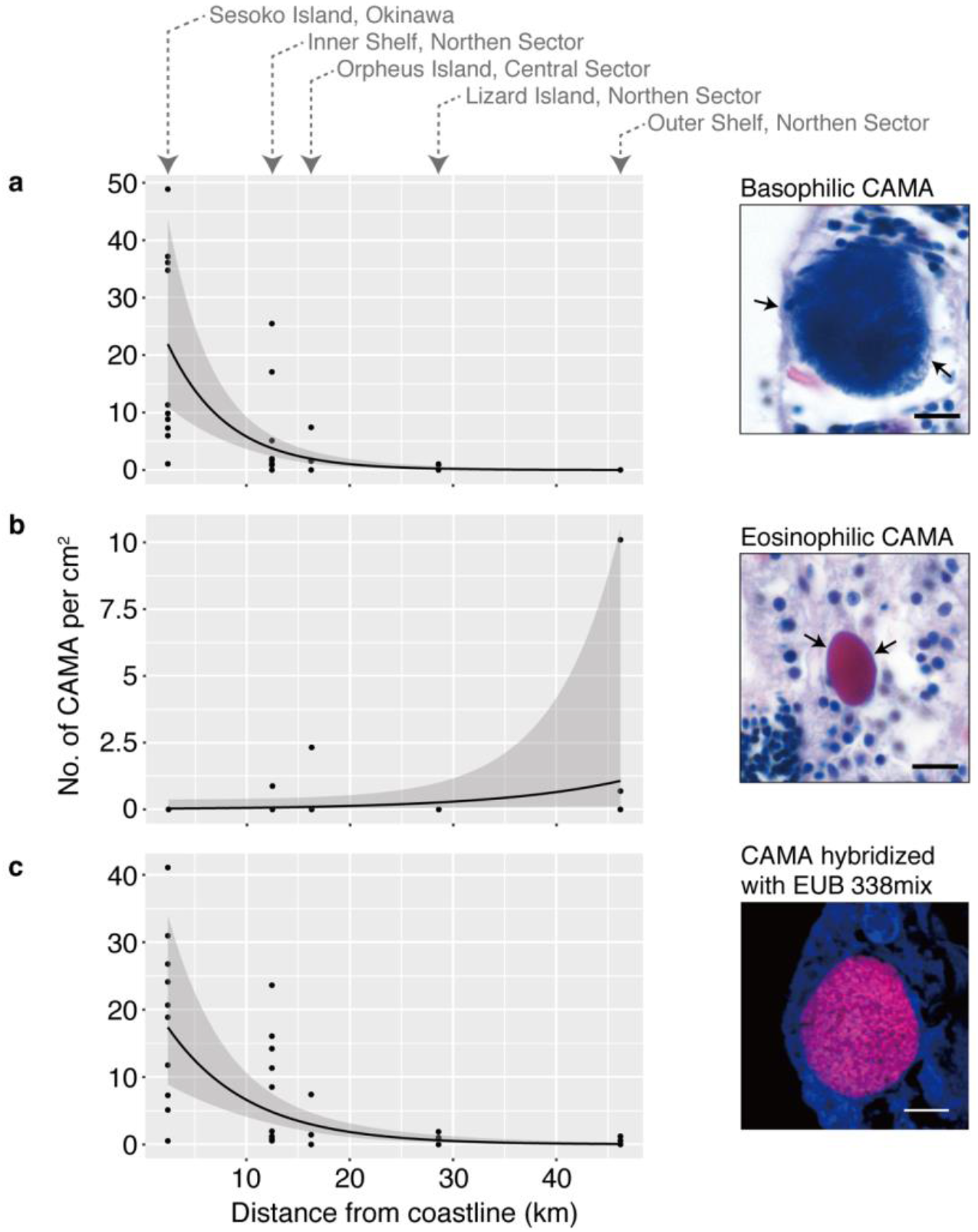
Relationship between densities of CAMAs and distance to the coast based on a negative binominal regression model. **(a)** Density of basophilic CAMAs was negatively correlated with the distance from the coast to the offshore sampling sites. **(b)** No correlation of eosinophilic CAMAs were detected in relation to the distance to the coast, although higher density (max. 10.1 cm^2^) was found in the Outer Shelf (offshore). **(c)** Density of the CAMAs detected by FISH was also negatively associated with the distance. Model prediction and associated 95% confidence intervals were obtained from a negative binominal regression model between the densities of CAMAs and the distance to the coastline **(a, c)**. Scale bars indicate 10 μm.

### Distribution of CAMAs within anatomical regions of the coral polyp

CAMAs were localized (by FISH) within the six anatomical regions of the coral polyp: the tentacle, actinopharynx, mesenterymesenterial filament, coenenchyme, and calicodermis (see **Fig. 4a**). The CAMAs appeared to be randomly distributed across the anatomical regions investigated, though because the numbers of CAMAs characterized for the Lizard Island, Outer Shelf and Orpheus Island samples were low (**Fig. 4b**), meaningful comparisons can only be made between the Sesoko Island and Northern Inner Shelf samples. For the Sesoko Island samples, CAMAs were found predominantly in the tentacles (36.5%), mesenterial filaments (34.7%) and coenenchyme (16.5%) regions; in Inner Shelf samples from Northern GBR sites, CAMAs were mostly located in the calicodermis (46.8%), tentacles (17.5%) and coenenchyme (16.7%) (**Fig. 4b**). An additional 31 CAMA-like shaped structures were observed from three samples derived from Sesoko Island, but these were discounted as bacterial aggregates due to non-probe binding signals and excluded from the analysis.

**Fig. 4.**
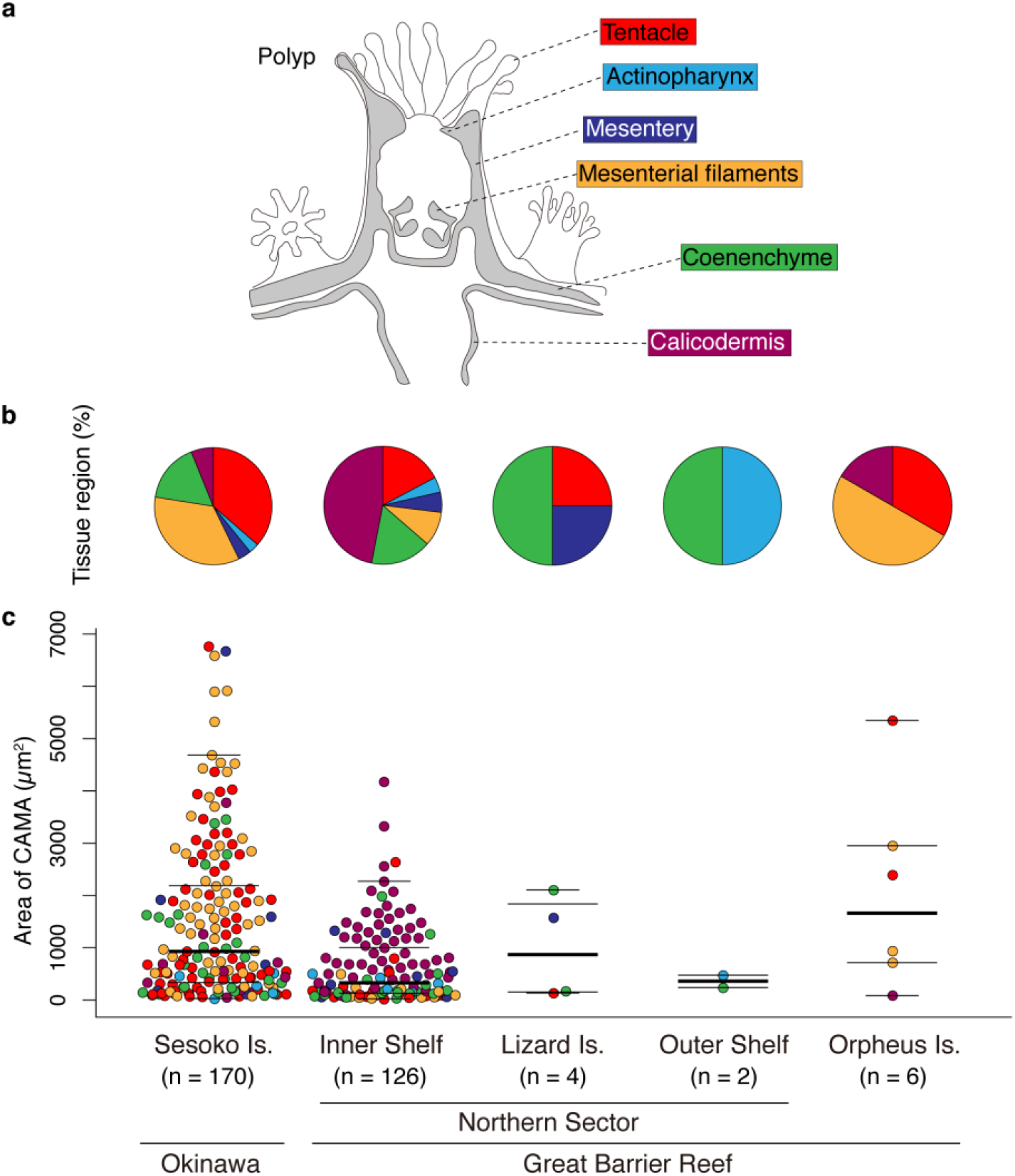
Distribution of CAMAs within six anatomical regions of the coral *Acropora hyacinthus* collected from five sites in Japan and Australia. **(a)** Schematic drawing of a coral polyp showing the six anatomical regions examined microscopically (same color coding used in **(b)** and **(c)**. **(b)** Pie charts showing the distribution of CAMAs among six anatomical regions in samples from the five sites. **(c)** Dot plots comparing the size of CAMAs among anatomical regions for each location. N=308 CAMAs detected in tissues treated with FISH. In the dot plot, the lines indicate median (thick line), 3^rd^ (upper) and 1^st^ (lower) quartiles (long thin lines), whiskers (short thin lines; upper value indicates largest value within 1.5* interquartile range from 3^rd^ quartile, lower: min value).

CAMAs spanned a wide size range, from 23 to 6,761 μm^2^ across tissue samples from all sites (**Fig. 4c**). Measurements likely underestimated the size of CAMAs, given they are dependent on the orientation of sectioning and it is unlikely that most CAMAs were sectioned through their greatest diameter. Acknowledging constraints associated with sectioning, the average size of CAMAs was 1,304 μm^2^. Sesoko Island samples contained generally larger average and median-sized aggregates (1,507.7±1,522.4 μm^2^, 921.3 μm^2^ respectively) than samples from the GBR region (Inner Shelf: 637.4±734 μm^2^, 321.6 μm^2^). However, given the large range in sizes measured, the fact that only two locations had sufficient sample sizes for comparison, the issue of sectioning orientation potentially biasing size measurements, and high variation among the individual colony level (**Suppl. fig. S1**), such patterns require further validation. No patterns in the size of CAMAs across different anatomical regions were detected (**Fig. 4c**).

### Morphology of CAMAs within coral tissues

High resolution imaging was used to partially characterize and compare the morphology of bacteria across the CAMAs detected (**Fig. 5**). Interestingly, each CAMA appeared to be composed of a single morphological type of bacteria, although morphological types varied among CAMAs. Overall, five different morphological types of bacteria were identified: rod-shaped (length 2.5±0.1 μm, width 0.6±0.0 μm, **Fig. 5a, e**), an atypical coccus (length 4.8±0.3 μm, width 3.1±0.2 μm, **Fig. 5b, f**), a longer rod morphology (length 8.0±0.0 μm, width 0.8±0.0 μm, **Fig. 5c, g**), filamentous-like bacteria (length N.D., width 0.4±0.0 μm, **Fig. 5d, h**), and a rod-shaped morphology but with spore-like structures (**Fig. 5i–j, m–n**). While the consistency of morphological characteristics within each CAMA may indicate that CAMAs are hosting single bacterial types, it is also possible that they host multiple bacterial species with similar morphologies. Interestingly, the fluorescent signal detected for some CAMAs was not uniform over the entire aggregation. The lack of signal within some CAMAs (see **Fig. 5d** for example) may be due to the probe not targeting microbial cells that inhabit that space. Defining a specific morphological shape for the bacterial cells observed was sometimes difficult, with patchy probe hybridization producing images of structures that potentially protruded from the tissue sections and seemed amorphous (**Fig. 5k–l, o–p**). However, this again could be the result of mixed microbial communities within the CAMAs, with some cells targeted by the probes, but others not hybridizing to the probe-fluorochrome conjugate.

**Fig. 5.**
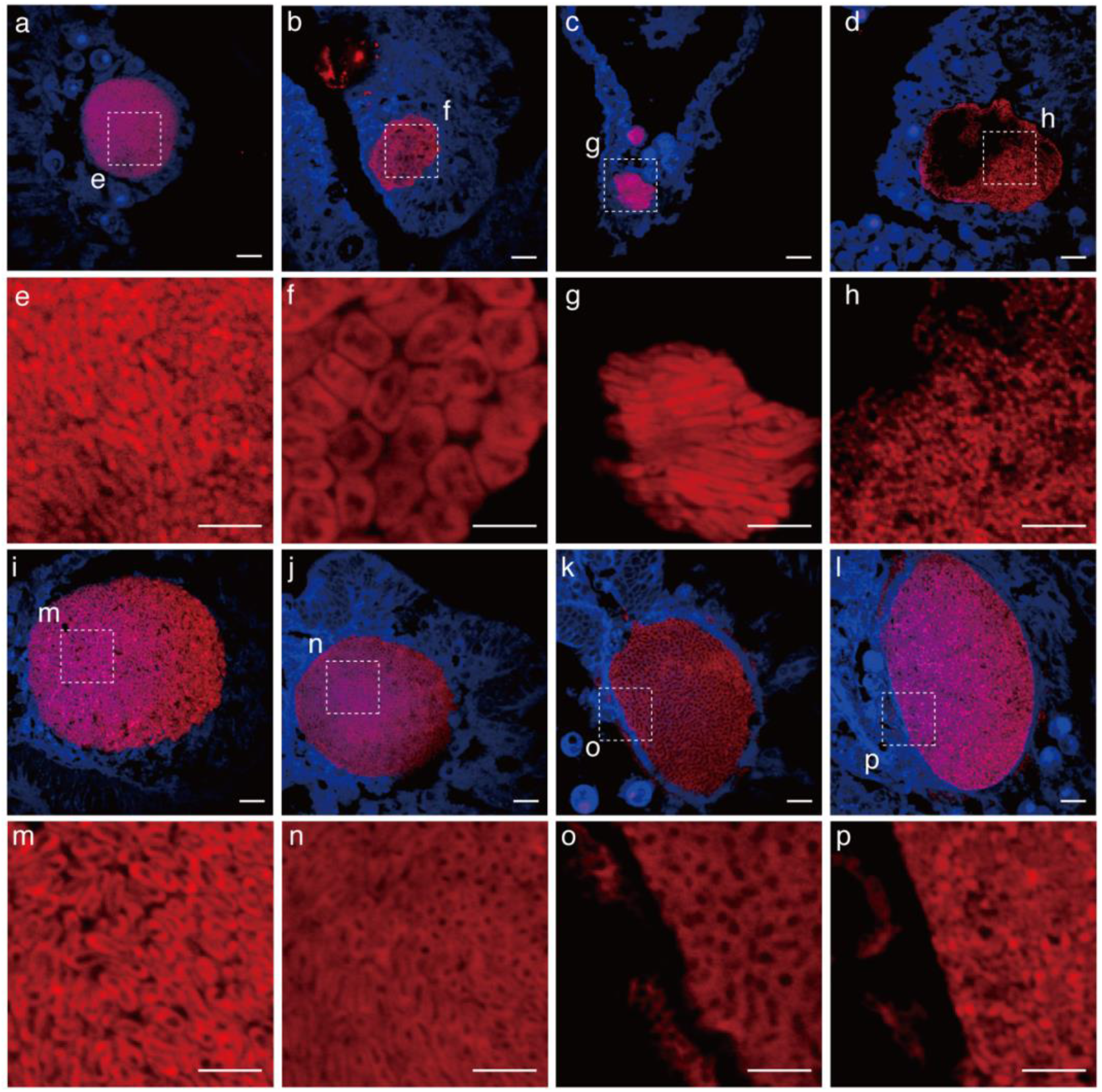
Morphological variation of bacteria housed within different CAMAs. Note that bacteria within each CAMA are morphologically similar. Dotted lines (in **a–d** and **i–l**) delineate regions magnified in close-up images (**e–h** and **m–p**). Overall, five bacterial morphologies were detected: rod-like (**a, e**), pleomorphic (**b, f**), long rods (**c, g**), filamentous-like (**d, h**), rod-shaped with spore-like structures (**i–j** and **m–n**), and putative amorphous masses (**k–l** and **o–p**). Scale bars indicate 10 μm (**a–d** and **i–l**) and 5 μm (**e–h** and **m–p**).

The variation in bacterial cell morphology among CAMAs may contribute to the different HE staining properties observed, with the majority being basophilic (95.9%), but 4.1% staining eosinophilic (n=318 CAMAs detected in total). Variability in the staining of CAMAs has been reported previously^32,51^ and attributed to varying degrees of protein and DNA production or local tissue pH conditions^32^. However, these staining patterns may also be influence by different bacteria housed within the CAMAs. Previous studies have reported that the abundant coral-associated bacterial genera *Endozoicomonas* forms aggregates within tissues of the corals *Stylophora pistillata* and *Pocillopora verrucosa*^38,39^. Indeed, extensive sequence-based phylogenetic surveys of coral microbiomes have revealed that several dominant bacterial groups are common^20^, including the *Proteobacteria* (particularly *Alpha*- and *Gammaproteobacteria* inclusive of *Endozoicomonas*), as well as *Actinobacteria, Bacteroidetes* (especially *Flavobacteria*), and *Cyanobacteria*^3^. Hence some of the different cellular morphologies we detected within CAMAs may represent common coral-associated bacterial groups profiled in microbiome diversity studies. We note, however, that bacterial morphology can be plastic and dependent on numerous variables, such as division stage, colonization, chemical environment, physical constraints and nutrient availability^52^. Future work targeting CAMAs with taxa-specific probes and gene-sequencing approaches would help to resolve both the taxonomy, composition and potential functional roles of bacteria within these structures of the coral holobiont.

FISH imaging showed that for some of the CAMAs, patchy probe-specific labeling occurred, resulting in dull or dark areas within the structures. There are a number of potential methodological reasons for this observation, including: 1) insufficient probe sensitivity due to low ribosomal rRNA content in target cells^53,54^, 2) methodological and environmental factors that prevent probes from accessing target cellular rRNA at these sites^55^, 3) the bacterial community penetrating and proliferating within the epidermis of coral tissues^37^, or 4) lipid or fat solvents deposited through dehydration and dewaxing steps showing empty spaces in the tissues^56^. Alternatively, little to no signal in central regions of some of the CAMAs could indicate that the probe EUB338mix did not target the taxonomic group of microbes within these regions. The EUB338mix is estimated to cover 96% of the *Eubacteria* domain, but taxa outside this coverage, including archaeal lineages, may be present^57^. We speculate that some of the CAMAs may be mixed communities containing bacteria not targeted by the probes or even *Archaea*, which have been identified to associate with corals in microbiome diversity studies^58^. A recent coral metagenomic study recovered *Thermarchaeota* genome bins from a *Porites* sp. that was potentially metabolically linked through nitrogen cycling to other coral microbial-associated taxa, including *Nitrospira*^6^. Co-aggregation of ammonia-oxidizing archaea with nitrite-oxidizing bacteria is common in other organisms, such as sponges^59,60^.

Further three-dimensional reconstructions of z-stacked FISH images of select 100 μm stained sections of tentacles visualized the CAMAs as typically spheroid or ellipsoid-shaped structures (**Fig. 6a**). In one example, a single CAMA was located independently in the epidermis of a tentacle (**Fig. 6b**). In a different tentacle, multiple smaller CAMAs were localized close to Symbiodiniaceae cells in the gastrodermis (**Fig. 6c**). Sizes of the large single CAMA and the multiple smaller CAMAs were 33,400 μm^3^ (**Fig. 6d**, surface area 7,729 μm^2^) and 1,978 ± 141.2 μm^3^ (**Fig. 6e**, 809.5 ± 59.9 μm^2^, n=4 CAMAs), respectively. Even though the CAMAs were located in the same polyp, the size of the single large CAMA was approximately 40-fold greater than the multiple smaller CAMAs, demonstrating inherent size variability for these structures. The bacteria within these CAMAs displayed a similar rod-shaped morphology (see **Fig.5a and e**), with the average cross-sectional area of each bacterium being 175.7 μm^2^ and 22.6 ± 4.2 μm^2^, respectively (**Suppl. fig. S2**). Based on cell size, the number of individual rod-shaped bacterial cells within the 3D rendered images of these CAMAs was estimated as ~ 47,275 cells for the large single CAMA located in the epithelium of the tentacle (**Fig. 6d** and **Suppl. fig. S2a**), and 2,799 ± 200 cells (**Fig. 6e** and **Suppl. fig. S2b**) for each of the smaller CAMAs localized close to Symbiodiniaceae cells.

**Fig. 6.**
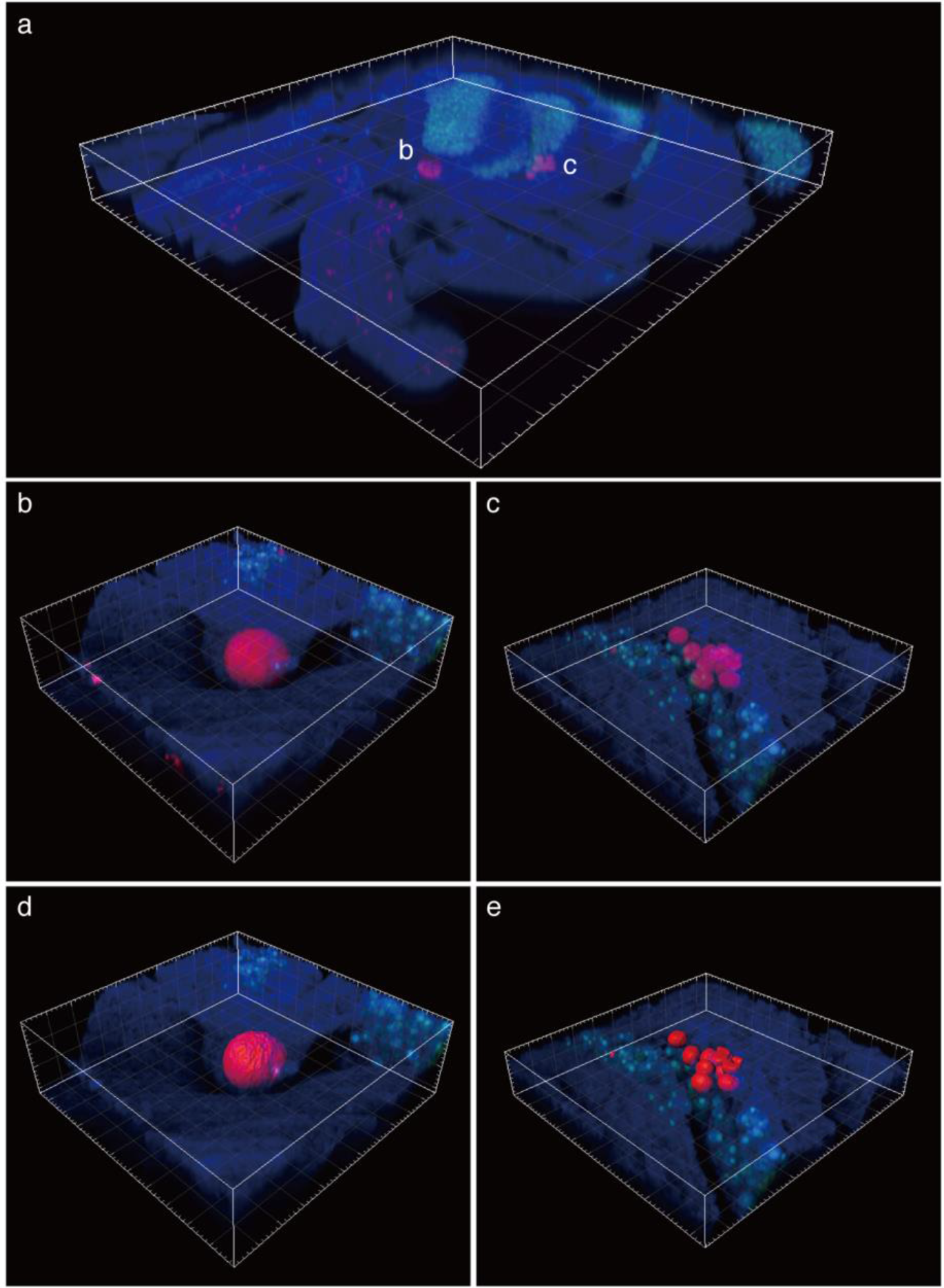
Three-dimensional (3D) images of CAMAs (red) within a tentacle of the coral *Acropora hyacinthus*, as visualized using FISH. **(a)** Section of tentacle showing localization of two types of CAMAs (10x magnification). **(b)** Single aggregation of bacteria in a large structure within the epithelium (40x magnification; composed of 92 z-stack images). **(c)** Multiple aggregations of bacteria in smaller structures within the gastrodermis (40x magnification; composed of 56 z-stack images). 3D rendering of CAMAs (**d, e**) reconstructed from 3D images in **b** and **c**. Coral tissue and Symbiodiniaceae appear as blue and green structures, respectively.

Using these estimates of cell numbers within CAMAs we further extrapolated that bacterial densities in tissues from Sesoko Island and Inner Shelf GBR corals are approximately 5.6 x 10^4^ and 1.5 x 10^4^ cells per cm^2^ (along a linear cross-section), respectively. Very few studies have accurately determined bacterial cell densities associated with corals. Counts for bacteria in the coral mucus layer can be as high as 10^7^ cells ml^−1^^61^. Estimates of bacteria on tissue surfaces range from 1 x 10^5^ to 10^6^ cells per cm^2^ for the coral *Pocillopora damicornis*^62^ and 8.3 x 10^6^ to 6.2 x 10^7^ cells per cm^2^ for the coral *Oculina patagonica*^24^. However, these counts are based on bacteria external to coral tissues and therefore not directly comparable to estimates from our 3D reconstructions of CAMAs visualized within coral tissues. Our results highlight that tissue-associated microbial communities are present across the different anatomical regions of the coral polyp and that differentiating these communities from external and mucus-associated microbial communities is important for accurate appraisals of the coral microbiome and for identifying their role(s) within the coral holobiont.

## Conclusion

Localization of microorganisms associated with corals is vital to understand their symbiotic relationships and reveal their function(s) within the holobiont. Here we provide novel insight into the distributions and densities of bacteria within tissues of the coral *Acropora hyacinthus* sampled from five different locations. CAMAs were common in coral tissues sampled, although their prevalence were higher in colonies sampled close to shore compared to offshore geographic sites, potentially linked to water quality parameters. While each CAMA appeared to be dominated by a single bacterial morphological type, different CAMA hosted different bacterial morphotypes. CAMAs have been defined as facultative symbionts, not necessary for host fitness^32^, however their high prevalence and abundance in coral tissues may indicate they are integrated into shared metabolic pathways and central to maintaining coral fitness through provisioning benefits. For example, findings that CAMAs are often co-localized near Symbiodiniaceae cells highlights the potential metabolic integrated links between bacterial and Symbiodiniaceae symbionts. Such propositions require testing by tracing metabolic pathways, as well as improved taxonomic and functional assessment of the microorganisms housed within these CAMAs.

## Methods

### Sample collections

Fragments (~ 3 cm x 3 cm; n=48 colonies) of the tabulate coral *Acropora hyacinthus* (Dana, 1846) were collected from five geographic locations across two countries (Australia and Japan). Samples from Japan (n=10) were collected from Sesoko Island, Okinawa (26°37’40.3”N, 127°51’36.4”E, depth 1.5–3.0 m, distance from coastline of Okinawa main Island 2.4 Km, average colony size 62.8 ± 41.7 cm) in July 2015. On the Great Barrier Reef (GBR) Australia, samples were collected from the Northern sector along an inshore to offshore gradient inclusive of Lizard Island (see **Fig. 1**; n=10 from an Inner Shelf site (14°47’29.4”S, 145°20’18.8”E, depth 3–5 m, distance from coastline 12.5 Km, average colony size 77.69 ± 43.0 cm), n=10 from Lizard Island (14°41’48.75”S, 145°27’49.22”E, depth 2–4 m, distance from coastline 28.6 Km, average colony size 61.67±17.5 cm) and n=8 from an Outer Shelf site (14°38’30.0”S, 145°38’22.5”E, depth 2–5 m, distance from coastline 46.2 Km, average colony size 41±20.5 cm) in January 2013. Samples (n =10) were also collected from reefs around Orpheus Island (18°35’55.4”S, 146°29’33.8”E, depth 2–3 m, distance from coastline 16.3 Km, colony size N.D.) in the inner-central region of the Great Barrier Reef in March 2013.

### Histological preparations

Following collection, samples were immediately rinsed with sterile seawater and then fixed in 4% paraformaldehyde (Electron Microscopy, USA and Wako, Japan) in 10mM phosphate buffered saline (PBS; pH 7.4) for 8–10 hours maintained at 4 °C.

Samples were subsequently rinsed twice with 70% ethanol and stored in 70% ethanol at 4 °C prior to decalcification. After samples were rinsed by PBS twice for 30 min each, samples were decalcified at 4 °C in a 10% EDTA (sigma-Aldrich, USA) solution (w/v; pH 8.0 adjusted by sodium hydroxide [Wako, Japan]), which was exchanged approximately every two days until no coral skeleton remained (approx. two weeks). The decalcified samples were rinsed in PBS, and dehydrated sequentially through 70%, 90%, and two changes in abs. 100% ethanol series (60 min each), then processed through a 1:1 solution of abs. 100% ethanol and toluene, and two changes in toluene (30 min each), and embedded in paraffin. Nine sections (three serial sections x three sets; interval = 100 μm between each set) of each coral fragment, each 4 μm thick, were cut from each paraffin-embedded sample (**Suppl. fig. S3**). In addition, thicker 100 μm sections were cut from coral tissues to allow reconstructions of three-dimensional configurations of the CAMAs (see below). Serial sections were mounted on one slide coated with egg-white glycerin and on two slides with poly-L-lysine solution (Sigma-Aldrich, USA) for hematoxylin and eosin (HE) staining and fluorescence *in situ* hybridization (FISH), respectively. We analyzed a total 144 sets (432 sections) of the serial sections for HE staining and FISH among five location sites.

### Hematoxylin and Eosin (HE) staining

For hematoxylin and eosin (HE) staining, one serial section from each set was dewaxed in xylene (2 x 15 min), rehydrated through ethanol series with abs. 100%, 99%, 90% and 70% (5 min each) and rehydrated completely in sterile water. Hydrated sections were stained in Mayer’s hematoxylin (Wako, Japan) for 10 min, rinsed in water for 5 min, then stained with eosin Y (Merck, Germany) for 5 min, and further rinsed in water for 30 sec. The stained sections were dehydrated through the same ethanol series in reverse with agitation (few sec each), cleared by xylene (2 x 5 min) and finally mounted in Entellan mounting medium (Merck, Germany). HE stained sections were observed and recorded using an ECLIPSE Ni microscope (Nikon, Japan) and BIOREVO BZ–9000 microscope (KEYENCE, Japan). All sampled corals appeared visually healthy at the time of collection and tissues displayed normal cell morphology, including no signs of fragmentation, wound repair or necrosis (as per criteria in^40^). The number of CAMA were counted into two categories, basophilic and eosinophilic CAMAs. The area of whole tissue (including skeleton region) was measured by tracing the outer edge of the tissue section using Fiji software^63^.

### Fluorescence in situ hybridization (FISH)

The other two serial sections from each set were subjected to FISH according to the protocol detailed in Wada et al. (2016)^64^. Briefly, sections were dewaxed in xylene (2 x 15 min), dehydrated briefly once in 100% ethanol and dried completely. The dried sections were immersed in a 0.2 M HCl solution for 12 min, followed by a 20 mM Tris-HCl solution (pH 8.0) for 10 min at room temperature. The sections were mounted with proteinase K (50 μg ml^−1^) in 20 mM Tris-HCl solution at 37 °C for 5 min for bacterial cell wall permeabilization, and washed in a 20 mM Tris-HCl solution at room temperature for 10 min. Oligonucleotide probes, including a probe targeting the 16S rRNA gene (EUB338mix: 5’-GCWGCCWCCCGTAGGWGT-3’) and a nonsense, negative control probe (Non338: 5’-ACATCCTACGGGAGGC-3’), were labeled with the Cy3 fluochrome (eurofins, USA)^65,66^. Tissue sections were covered with hybridization buffer (30% v/v formamide, 0.9 M NaCl, 20 mM Tris-HCl [pH 8.0], 0.01% SDS), then each oligonucleotide probe was added to a final concentration of 25 ng μl^−1^ to each serial section. The slides were incubated at 46 °C for 1.5 hour. After incubation, sections were washed in 50-ml falcon tubes containing preheated wash buffer (0.112 M NaCl, 20 mM Tris HCl [pH 8.0], 0.01% SDS, 5 mM EDTA [pH 8.0]) in a water bath at 48 °C for 10 min, then soaked immediately with agitation in cold water, and air dried completely. The dried sections were mounted in an antifade mounting medium Fluoromount/Plus (Diagnostic BioSystems, USA). Sections were examined and recorded using a FV1000-D confocal microscope (Olympus, Japan) with two channels, using the following settings: (1) laser: 405 nm and 559 nm; (2) excitation dichroic mirror: DM405/473/559; (3) emission dichroic mirror: SDM560 and mirror; (4) band-pass filter: None and BA575–620 for detecting autofluorescence of coral tissue (blue) and Cy3 signal (red), respectively. Non-specific probe binding in tissue sections was identified as detailed by Wada et al. (2016)^64^. Specific detection of bacteria by fluorescent *in situ* hybridization (FISH) can be problematic for coral samples due to nonspecific binding of probes and background autoflorescence of granular cells and nematocysts^64^. Therefore, CAMAs were identified by their characteristic shapes, in addition to fluorescent signals bound by the general bacteria-targeted probe set EUB338mix and comparisons to background autofluorescence and non-specific binding (using Non338 probe). The number of CAMAs that hybridized with the EUB338mix probe in each section was counted and the area of whole tissue (including skeleton region) was measured by tracing the outer edge of the tissue section. We also traced the edges of each CAMA to calculate the area (the size of CAMA was dependent on the orientation of sectioning). Image processing was conducted in Fiji software^63^.

### Reconstruction of three-dimensional images of CAMAs

Three-dimensional images (3D) of CAMAs were reconstructed from the thick sections (100 μm) which were cut carefully from two samples (S6 from the Sesoko Island and LO8 from the Outer Shelf GBR site). Sections were visualized via FISH according to methods described above, and examined using a LSM 880 (ZEISS, Germany) with two tracks and the following settings: (1) laser: 405 nm and 561 nm; (2) beam splitter: MBS-405 and MBS 488/561; (3) filter: 371–479 nm for auto-fluorescence of coral tissue (blue) and 627–758 nm for Symbiodiniaceae (green) in track 1 and 565–588 nm for Cy3 signals (red) in track 2, respectively. For reconstructing the 3D images, the sections consisted of z-stack images at 3.0 μm intervals for 10x and 0.7 μm each and 40x magnifications using the Z-stack function in LSM 880. The z-stack images were processed and reconstructed with surface rending of the Cy3 signals in Imaris software ver. 8.0.2 (BitplaneAG, USA).

### Statistical analysis

All statistical analyses were conducted using R Stats ver. 3.5.1^67^ with the following packages: rcompanion ver. 2.0.0^68^, MASS 7.3-51.4^69^, pscl 1.5.2^70^ and car 3.0-2^71^. To compare the abundances of CAMAs at the colony level among sites, Fisher’s exact test followed by post hoc Benjamini–Hochberg false discovery rate correction were used. A generalized linear model was used to evaluate the relationship between the distance of samples from the coastline (explanatory variable) with the distribution and density of CAMAs (count data of response variables: basophilic, eosinophilic and FISH detected CAMAs; offset variable: tissue area) among the five sites. The tests were performed to fit using the ‘glm.nb’ function (negative binomial distribution) in the MASS package, due to over dispersions among the data. We also performed zero-inflated negative binomial regression using the ‘zeroinf’ function in the pscl package, since the count data included excessive zeros. We then performed a Vuong test that compares an ordinary negative binomial regression model against the zero-inflated model using the ‘vuong’ function in the pscl package, and adopted the model’s fitness based on AIC (Akaike’s Information Criterion), and yielded the model, as appropriate. Finally, the likelihood-ratio (LR) chi-square was calculated for each model using the car package.

## Supporting information

all supplemental informations

## Author contribution

NW, NM and DGB conceived the study. NW, FJP and DGB designed the sampling and FISH experiments. NW and FJP conducted the field sampling. NW, MI and TM performed the histological work and all observations and data collection. NW and DGB had major contribution in the manuscript writing and the figure making. FJP, S-LT, TDA, BLW and NM contributed to writing and editing the manuscript. All authors critically reviewed, revised and ultimately approved this final version. Finally the authors confirm that there are no competing interests and no non-financial competing interests that impact this manuscript

## Acknowledgements

Dr. Joleah Lamb (University of California, Irvine), Mr. Sefano Katz (Pacific Blue Foundation), Dr. Gergely Torda (ARC Center of Excellence for Coral Reef Studies, James Cook University), and Dr. Melanie Trapon (CSIRO) are thanked for helping to collect samples. We thank A/Prof Esther Peters from George Mason University for her advice and help in updating coral anatomy terminology. We are also grateful to Mr. Yasuhiko Sato from Zeiss Microscopy Japan, and Mr. Ji-Ying Huang and IPMB Live-Cell-Imaging Core Lab in the Institute Academia Sinica for supporting the microscopy and the Imaris software used.

