## Supplementary material for "Characterization of coral-associated microbial aggregates (CAMAs) within tissues of the coral *Acropora hyacinthus*": all supplemental informations

**Suppl. table S1** Fisher's exact test values for the presence/absence of CAMA detected by HE stain and FISH among samples collected from the five sites

| | Compared element | p-value of Adj. Fisher exact test |
|---|---|---|
| CAMAs detected by HE† | Sesoko Is. vs. Inner Shelf | 0.6770 |
| | Sesoko Is. vs. Lizard Is. | 0.0155** |
| | Sesoko Is. vs. Outer Shelf | 0.0151** |
| | Sesoko Is. vs. Orpheus Is. | 0.0360** |
| | Inner Shelf vs. Lizard Is | 0.1400 |
| | Inner Shelf vs. Outer Shelf | 0.1340 |
| | Inner Shelf vs. Orpheus Is. | 0.2830 |
| | Lizard Is. vs. Outer Shelf | 1.0000 |
| | Lizard Is. vs. Orpheus Is. | 1.0000 |
| | Outer Shelf vs. Orpheus Is. | 0.7980 |
| CAMAs detected by FISH | Sesoko Is. vs. Inner Shelf | 1.00000 |
| | Sesoko Is. vs. Lizard Is. | 0.00517*** |
| | Sesoko Is. vs. Outer Shelf | 0.00378*** |
| | Sesoko Is. vs. Orpheus Is. | 0.00357*** |
| | Inner Shelf vs. Lizard Is | 0.00517*** |
| | Inner Shelf vs. Outer Shelf | 0.00378*** |
| | Inner Shelf vs. Orpheus Is. | 0.00357*** |
| | Lizard Is. vs. Outer Shelf | 1.00000 |
| | Lizard Is. vs. Orpheus Is. | 1.00000 |
| | Outer Shelf vs. Orpheus Is. | 1.00000 |

†The HE stained CAMAs include basophilic and eosinophilic CAMAs.
P values are $**p < 0.05$ and $***p < 0.01$.

**Suppl. table S2** Summary of the densities of CAMAs in HE- and FISH- stained tissues

| | Sesoko Is. | Inner Shelf | Lizard Is. | Outer Shelf | Orpheus Is. |
|---|---|---|---|---|---|
| Sample size | 10 | 10 | 10 | 8 | 10 |
| **Basophilic CAMAs** | | | | | |
| Colonies observed with CAMAs | 10 | 8 | 3 | 0 | 3 |
| Ave. densities (n/cm$^2$) in tissues | 20.13±17.1 | 6.78±9.3 | 0.92±0.1 | - | 3.55±3.3 |
| Density range (n/cm$^2$) across samples | 1.06 - 48.90 | 0.86 - 25.48 | 0.87 - 1.02 | - | 1.54 - 7.42 |
| **Eosinophilic CAMAs** | | | | | |
| Colonies observed with CAMAs | 0 | 1 | 0 | 2 | 1 |
| Ave. densities (n/cm$^2$) in tissues | - | 0.86 | - | 5.39±6.7 | 1.28±2.8 |
| Density range (n/cm$^2$) across samples | - | - | - | 0.69 - 10.10 | |
| **FISH detected CAMAs** | | | | | |
| Colonies observed with CAMAs | 10 | 10 | 3 | 2 | 2 |
| Ave. densities (n/cm$^2$) in tissues | 18.72±12.7 | 7.90±8.2 | 1.26±0.6 | 0.90±0.5 | 4.48±4.2 |
| Density ranges (n/cm$^2$) across samples | 0.53 - 41.08 | 0.86 - 25.48 | 0.87 - 1.02 | 0.58 - 1.22 | 1.54 - 7.42 |

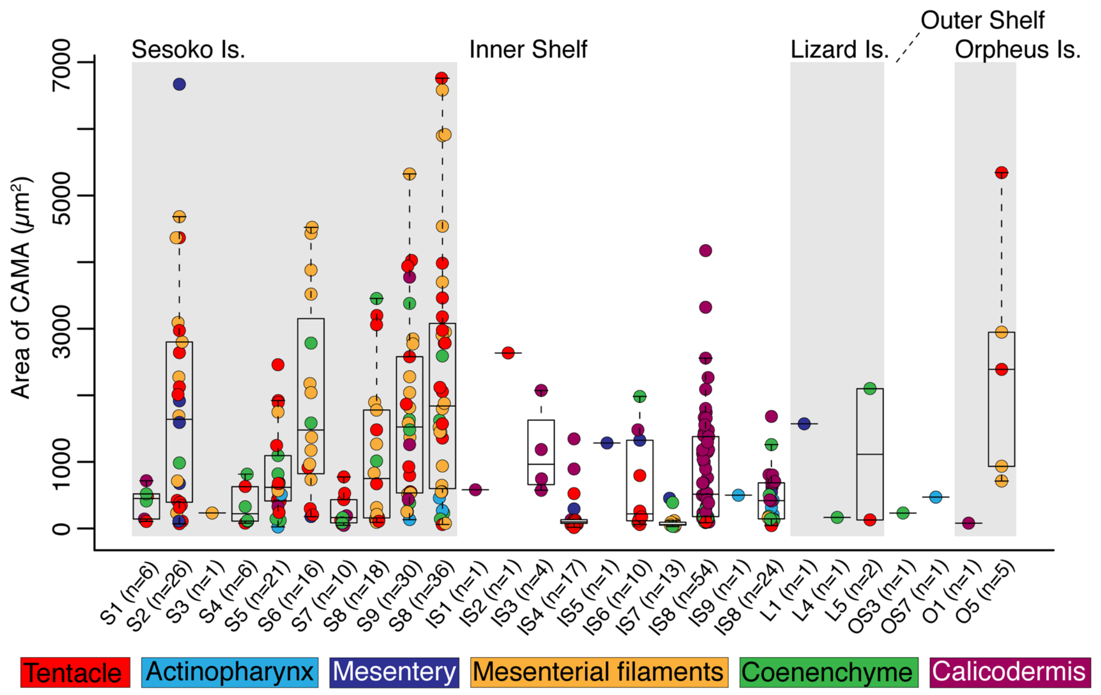


**Suppl. Fig. S1 High variation of CAMA size at the individual colony level.** Dot plots with box plots showing the size of CAMAs among six anatomical regions (see color coding in below index) for each site.

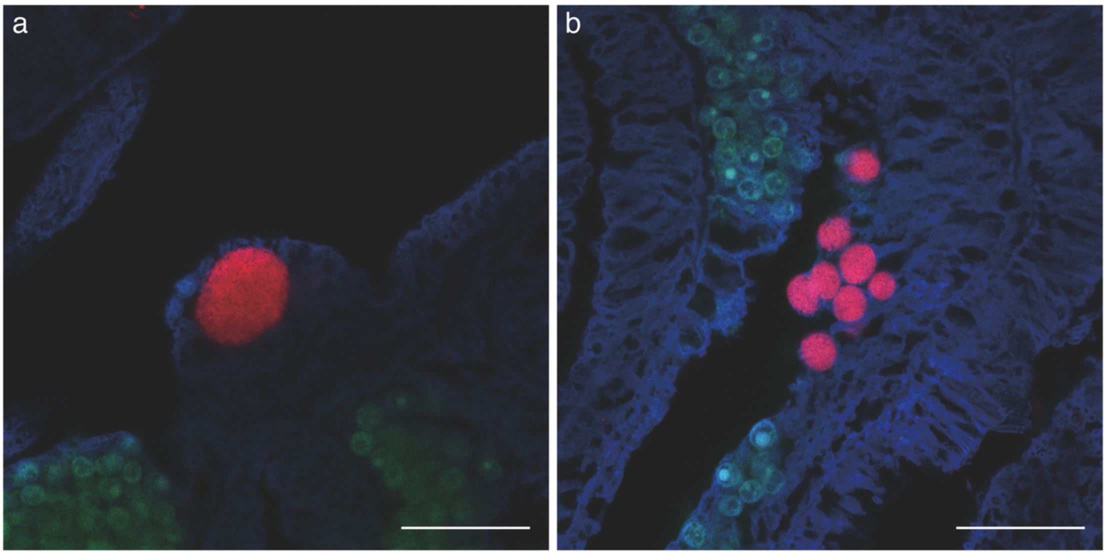


**Suppl. Fig. S2 Calculating the area of CAMAs. (a)** A single large CAMA, comprised of rod-shaped bacteria, was calculated to be 175.7 $\mu m^2$ in cross-sectional area from 3D images. **(b)** Smaller, numerous aggregations were calculated to be, on average, 22.6±4.2 $\mu m^2$ in cross-sectional area (n = 8). Scale bars indicate 50 µm.

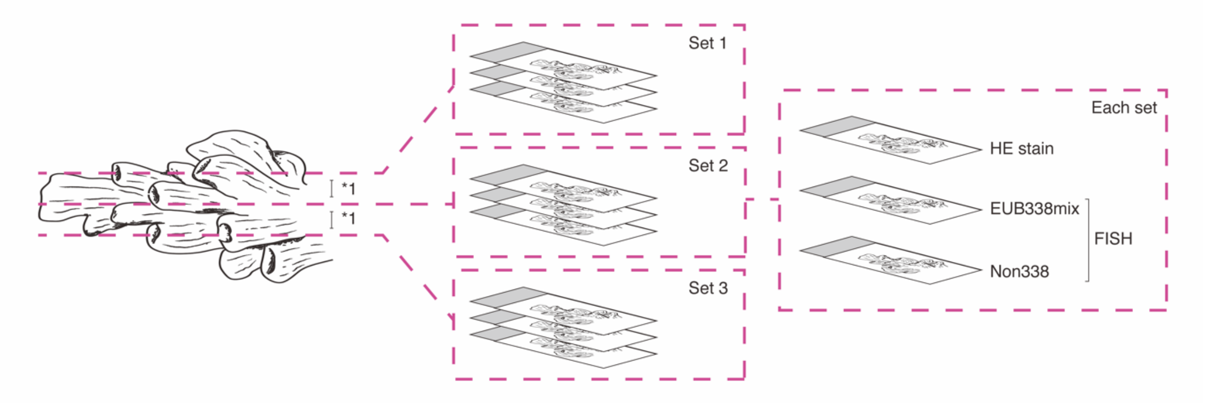


**Suppl. Fig. S3 Schematic drawing showing how coral fragments were sectioned for HE staining and FISH.** In total, nine sections were collected from each sample (three sets of sections, each set comprised of three serial sections). *1: Distance between each set was 100 µm.
